# A whole-genome screen identifies a bilateral Flower-Grindelwald/TNFR-Veli/LIN-7 communication module in cell competition

**DOI:** 10.64898/2026.08.14.743262

**Authors:** Catarina Marçal-Costa, Catarina Brás-Pereira, Dina S. Coelho, Sara Astrid Vilão, Marta Curado-Avelar, Joana Couceiro, Christa Rhiner, Eduardo Moreno

## Abstract

Viable yet suboptimal (“loser”) cells can be recognized and selectively eliminated when in the presence of neighboring fitter (“winner”) cells through cell competition, thereby promoting optimal tissue fitness and homeostasis. One mechanism through which cells compare their relative fitness levels relies on isoforms of the conserved transmembrane protein Flower (Fwe). Despite the role of Fwe-dependent cell selection in several pathophysiological processes, little is known about downstream components of this pathway. In this study, we develop a versatile clonal interaction assay in *Drosophila*, the Easy Win Assay. By overexpressing human FWE1 Lose isoform to trigger cell competition, we perform an unbiased whole-genome RNAi screen and identify new pathway modulators. We show that the receptor Grindelwald/TNFR is necessary in either winner or loser cells to drive Eiger/TNF-α-independent elimination of losers, whereas the scaffolding protein Veli/LIN-7 is simultaneously required in both cell populations. We further demonstrate that Fwe, Grindelwald and Veli redistribute to establish a previously undescribed bilateral communication module at winner-loser interfaces, promoting intercellular communication and elimination of loser cells. These findings show that distinct cell elimination pathways converge on a common execution module while remaining independently regulated upstream, enabling flexible recognition and elimination of diverse damaged or dangerous cells.

## Introduction

Multicellular organisms rely on continuous cell-cell communication to coordinate organ function and physiological mechanisms, thereby maintaining tissue homeostasis (Su et al., 2024). During development and adulthood, viable yet suboptimal cells can arise within a tissue due to exposure to external insults or disrupted internal cellular processes (Baker, 2020; Khandekar & Ellis, 2024). To optimize tissue fitness and prevent the detrimental accumulation of such unfit cells over time (Akagi et al., 2018; Merino et al., 2015), these “loser” cells are recognized and eliminated when surrounded by fitter neighboring “winner” cells, which expand at their expense, via a Darwinian-like selection termed cell competition (Morata, 2021; Moreno & Rhiner, 2014). This context-dependent, non-cell-autonomous process relies on cell-cell interactions and on relative, rather than absolute, cell fitness levels (Costa et al., 2020; Vivarelli et al., 2012), and has emerged as a fundamental mechanism to optimize tissue fitness with important implications for development, neurodegeneration and tumorigenesis (Baker, 2020; Kim & Jain, 2020; Morata, 2021). Yet, how fitness information is read between neighboring cells and converted into selective elimination of losers is not fully understood.

One mechanism through which cells compare their relative fitness relies on “fitness fingerprints” and direct cell-cell contact (Madan et al., 2018; Merino et al., 2016; Moreno & Rhiner, 2014). Cells present different isoforms of the conserved transmembrane protein Flower (Fwe), labeling them according to their fitness status. In *Drosophila*, loser cells express Fwe Lose isoforms, while winner cells express Fwe Ubi, which is slightly downregulated in losers (Rhiner et al., 2010). In parallel, transiently unfit cells upregulate *SPARC*, which encodes a secreted protective protein that elevates the threshold for Caspase activation (Portela et al., 2010). However, if a cell irreversibly acquires a loser fate and shares 40-50% of its surface with winners (Levayer et al., 2015), *ahuizotl* (*azot*) expression is activated. Azot integrates these different fitness cues to drive *head involution defective* (*hid*) upregulation, triggering apoptotic elimination of losers (Merino et al., 2015). The *fwe* gene is functionally conserved across species (Madan et al., 2019; Petrova et al., 2012), and human isoforms can activate competition in *Drosophila* tissues, with human Flower 1 (hFWE1) acting as a Lose isoform (Costa-Rodrigues et al., 2025). Fwe-dependent cell selection has been shown to influence several processes, including development, tissue repair, lifespan and Alzheimer’s disease (Coelho et al., 2018; Coelho & Moreno, 2020; Li et al., 2024; Merino, 2023; Merino et al., 2013, 2015; Moreno et al., 2015). Additionally, if hijacked by premalignant cells, it can promote tumorigenesis. Genetic or therapeutic targeting of this mechanism in mammals has been shown to reduce carcinogenesis and sensitize malignant cells to chemotherapy (Madan et al., 2019, 2025; Petrova et al., 2012). Despite the physiological and pathological relevance of this cell fitness-driven pathway, it remains poorly understood. Key open questions include identifying the molecular partners that synergize with Fwe as fitness sensors and defining the active role of winner cells during competition. Performing an unbiased genome-wide genetic screen represents a classic and effective strategy to address these gaps.

In this study, we developed a novel genetic design, the Easy Win Assay (EWA), a versatile clonal interaction assay that enables the targeted manipulation of defined cell populations to perform a genome-wide RNAi screen in *Drosophila*. Through this approach, we obtained a diverse set of enhancers and suppressors of Fwe-triggered clonal elimination and identified Grindelwald (Grnd)/TNFR and Veli/LIN-7 (Andersen et al., 2015) as pivotal determinants of Fwe-dependent clonal competition, uncovering previously unrecognized roles for these interactors in tissue surveillance. Together with Fwe, Grnd and Veli function as a module at the winner-loser interface to promote loser cell elimination and maintain epithelial homeostasis. Ultimately, we propose that the Fwe-Grnd-Veli module acts as a bilateral mediator of fitness-driven intercellular communication.

## Results

### Clonal interactions revealed by the “Easy Win Assay”

Genetic manipulation of specific cell populations is essential for advancing our understanding of cell-cell communication. With the goal of screening for new candidate genes involved in Fwe-dependent cell competition in a high-throughput manner, we developed a functional genetic assay in the *Drosophila* eye, referred to here as “Easy Win Assay” (EWA). The versatile genetic system is adaptable to study clonal interactions and allows for straightforward scoring of functional outcomes by examining the resulting proportion of differently marked clones in the adult eye tissue, which is used as a readout (**Figure 1A,B**). Genetically, a flippase recombinase (FLP) is driven by the *eyeless* (*ey*) promoter, which is active in the eye-antennal disc primordium during embryonic and first-instar (L1) larval stages, becomes restricted to the eye domain in L2, and is turned off posterior to the morphogenetic furrow upon cell differentiation. To generate clones, we created a *CoinFLP LHV2* cassette that allows the formation of LHV2-positive clones, using a refined version of the LexA transactivator that achieves higher expression levels while avoiding adverse developmental side effects (Bosch et al., 2015; Yagi et al., 2010). Stochastic FLP/FRT recombination leads to either the excision of STOP sequence, enabling clonal LHV2 expression (*act-FRT-FRT3-LHV2*), or permanent STOP maintenance (*act-FRT-STOP-FRT3*), in approximately equal cell populations (**Figure 1A**). Activation of *LexAop* (*Lop*) sequences by LHV2 drives co-expression of transgenes (*Lop-Y*) and *white RNAi* (*Lop-white RNAi*). This enables the visual tracking of LHV2-expressing clones as unpigmented (white) tissue within an otherwise pigmented fly eye (**Figure 1A,B**). To manipulate both the clones and the remaining cells, Gal4 is broadly expressed in the *ey* domain through highly efficient flip-out of FRT-flanked elements in the integrated *actin-stop-Gal4* construct (*act5-FRT-STOP-FRT-Gal4*) (**Figure S1A,B**), thus resulting in overexpression (*UAS-gene X*) or knockdown (*UAS-gene X RNAi)* of genes of interest in both cell populations. The additional inclusion of *UAS-Dicer2* enhances RNAi efficiency.

**Fig. 1.**
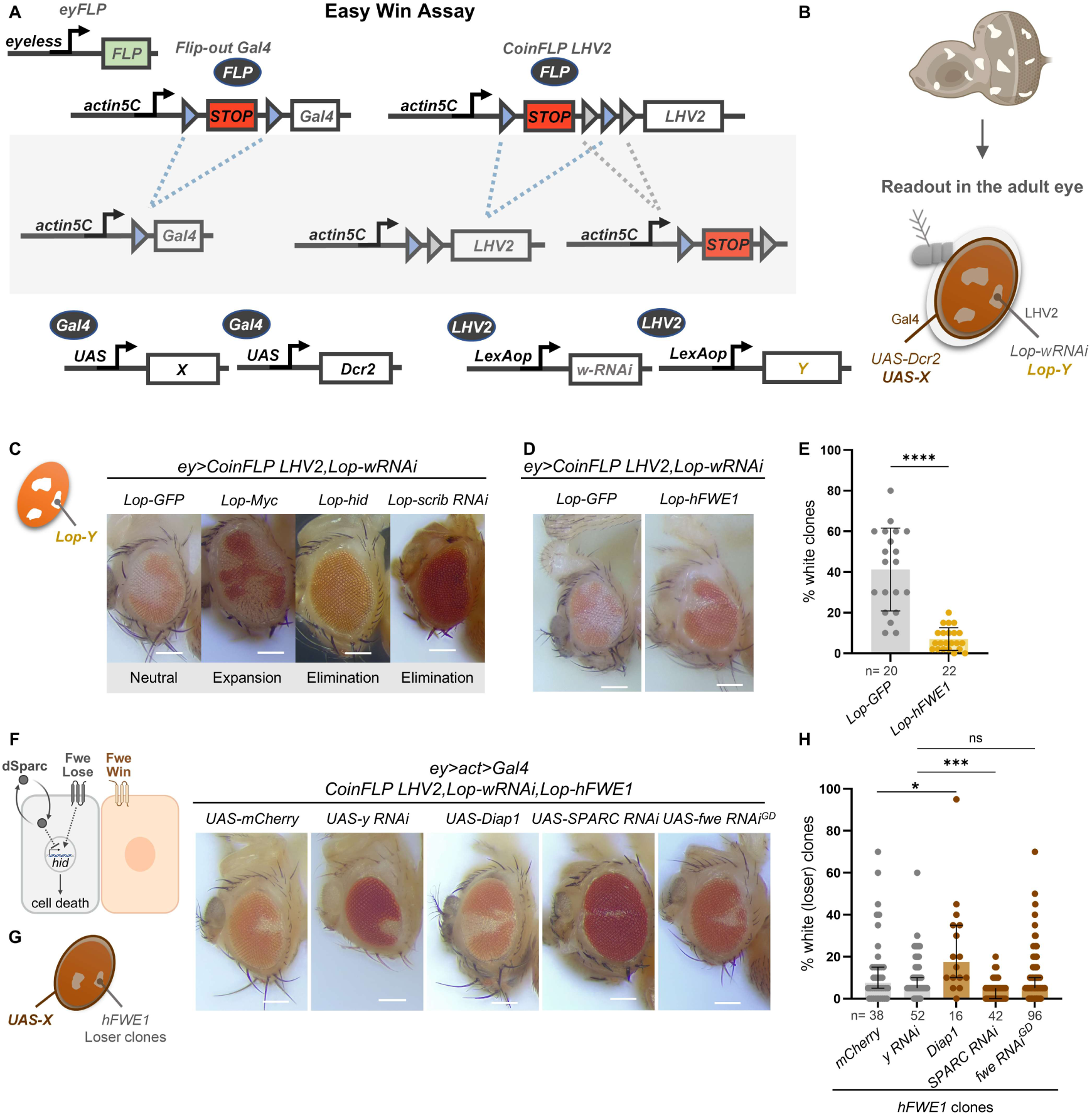
Easy Win Assay (EWA) – a tool to easily visualize the outcome of clonal interactions in the adult eye. **A.** *Eyeless* (*ey*) flippase (FLP) acts on FRT (blue triangles) and FTR3 (grey triangles) sites, present on *flip-out Gal4* and *CoinFLP LHV2* cassettes. Two distinct binary expression systems are present – *Gal4/UAS* and *LHV2/LexAop* (*Lop*). Different *UAS-X* and *Lop-Y* can be used to personalize the EWA tool. **B.** Active clonal interactions occur in the *ey* domain and the adult eye is used as a readout to phenotypically assess the resulting outcome. The *LHV2/Lop* clonal subset is labeled in white (*Lop-wRNAi*), and the *Gal4/UAS* system is active in both cell populations. **C.** Representative images of the resulting outcome of clonal overexpression (*ey>CoinFLP LHV2,Lop-wRNAi,Lop-Y*) of control *GFP*, *Myc*, *hid* and *scrib RNAi*, labeled as white tissue. **D.** Clonal overexpression of *GFP* or *hFWE1* labeled in white (*ey>CoinFLP LHV2,Lop-wRNAi,Lop-Y*). **E.** Quantification of the percentage of white clones in relation to the total eye size. Crosses at 29°C; n=number of eyes (females); Unpaired t test. **F.** Loser cells present the Fwe Lose isoform at the membrane, while winner cells present Fwe Win (Fwe Ubi). Losers express the secreted protein dSparc but ultimately trigger the activation of downstream cell death effectors. **G.** Representative images of adult eyes containing *hFWE1* white loser clones and overexpression of different transgenes in the entire eye (*ey>CoinFLP LHV2,Lop-hFWE1,Lop-wRNAi; >act>Gal4,UAS-X*). **H.** Quantification of the percentage of recovered white loser tissue in each condition. Crosses at 29°C; n=number of eyes (females); Mann-Whitney test and Kruskal-Wallis test (multiple comparisons).

The combination of two expression systems, *Gal4/UAS* and *LHV2/Lop*, and the choice of different *UAS-X* and *Lop-Y* elements enables flexible and precise control over the EWA. To demonstrate this versatility and validate the tool’s functionality, we overexpressed transgenes known to promote clonal expansion such as *Myc* (De La Cova et al., 2004; Moreno & Basler, 2004), or clonal elimination such as overexpression of the pro-apoptotic gene *head involution defective* (*hid*) (Grether et al., 1995) or downregulation of the cell polarity gene *scribble* (*scrib*) (Brumby & Richardson, 2003; Igaki et al., 2009). The outcome of the interaction between the manipulated *LHV2/Lop* clonal subset *versus* the *LHV2*-negative population was directly assessed by analyzing the proportion of white tissue in the adult eye in comparison to a neutral control (*Lop-GFP*). As expected, we detected an increased amount of white tissue for *Myc*-overexpression (*Lop-Myc*) and elimination of white areas upon activation of *hid* (*Lop-hid*) or the knockdown of *scrib* (*Lop-scrib RNAi*) (**Figure 1C**). These findings demonstrated the capacity of the EWA to reliably report both clonal expansion and elimination.

### hFWE1 clones are eliminated through cell competition

Next, we sought to harness the EWA to explore Fwe-dependent cell competition. We customized the EWA overexpressing human Flower 1 (*Lop-hFWE1*) in clones, which confers loser status and acts as a potent trigger for cell competition-induced cell elimination in flies. The hFWE1 isoform functions analogous to *Drosophila fwe Lose B*, as previously shown (Costa-Rodrigues et al., 2025), therefore representing a humanized model. As expected, clones overexpressing *hFWE1* are eliminated from eye tissue (**Figure 1D,E**). Accordingly, analyzing *hFWE1* clones (*UAS-hFWE1*) in the wing imaginal disc using *hs-FLP*-mediated induction of *CoinFLP Gal4* cassette revealed that clone elimination is not a tissue-specific phenomenon (**Figure S1D,E**). In turn, uniform *hFWE1* overexpression (*tub>hFWE1*) did not trigger elimination (**Figure S1F**) – a hallmark of cell competition (Rhiner et al., 2010; Vivarelli et al., 2012). Interestingly, clonal *hFWE1* overexpression produced rough eye tissue in contrast to normal eye appearance when expressed uniformly (**Figure 1D and Fig S1F,G**). This phenotype suggests that clone elimination in the developing eye disc interferes with eye tissue differentiation and/or organization. Tissue-wide overexpression of *Death-associated inhibitor of apoptosis 1* (*Diap1*) rescued *hFWE1* loser clones from elimination, in contrast to overexpression of control *mCherry*, confirming that cell elimination is apoptosis-dependent. Conversely, downregulation of *SPARC*, a Fwe Lose antagonist that transiently protects loser cells from elimination in flies (Portela et al., 2010), further increased *hFWE1* clones’ elimination (**Figure 1F-H**). Last, we found that targeting of *Drosophila fwe* isoforms does not interfere with hFWE1-mediated cell elimination (**Figure 1F,H**), indicating that hFWE1 activation is sufficient to impose the loser state. Overall, these results show that, in the EWA, *hFWE1* clones behave as losers and their elimination involves known players of the Fwe pathway.

Additionally, we observed sexual dimorphism in the EWA setting. Male flies consistently presented fewer clones than females (**Figure S1H**). We also tested different temperatures and found that 29°C yielded the most robust readouts for clone elimination. Thus, we chose the optimized conditions (29°C; F1 females for scoring, except for RNAi lines on Y) for the subsequent screening for suppressors and enhancers of Fwe-dependent elimination.

### A whole-genome RNAi screen to identify modulators of hFWE1-induced cell competition

To uncover novel regulators of the Fwe pathway, we set out to perform a genome-wide RNAi screen to identify suppressors and enhancers of hFWE1-triggered cell elimination, which can reveal downstream and parallel effectors of the pathway. To find modifiers, we crossed EWA hFWE1 flies with one RNAi line *per* gene from the VDRC stock collection, which covers approximately the whole genome (12 508 protein coding genes). RNAi leads to the downregulation of target genes in the screened progeny, both in *hFWE1* loser clones and surrounding winner cells (**Figure 2A,B**). We preferentially tested RNAi lines from the KK collection, resorting to GD and shRNA RNAi lines when no KK line was available. As approximately 25% of lines of the KK collection may carry an extra insertion at *40D* driving ectopic *tiptop* expression (Vissers et al., 2016), we confirmed that it did not affect the elimination of *hFWE1* clones (**Figure S2A,B**). However, this insertion produced an unexpected side effect on antennal development (**Figure S2A**).

**Fig. 2.**
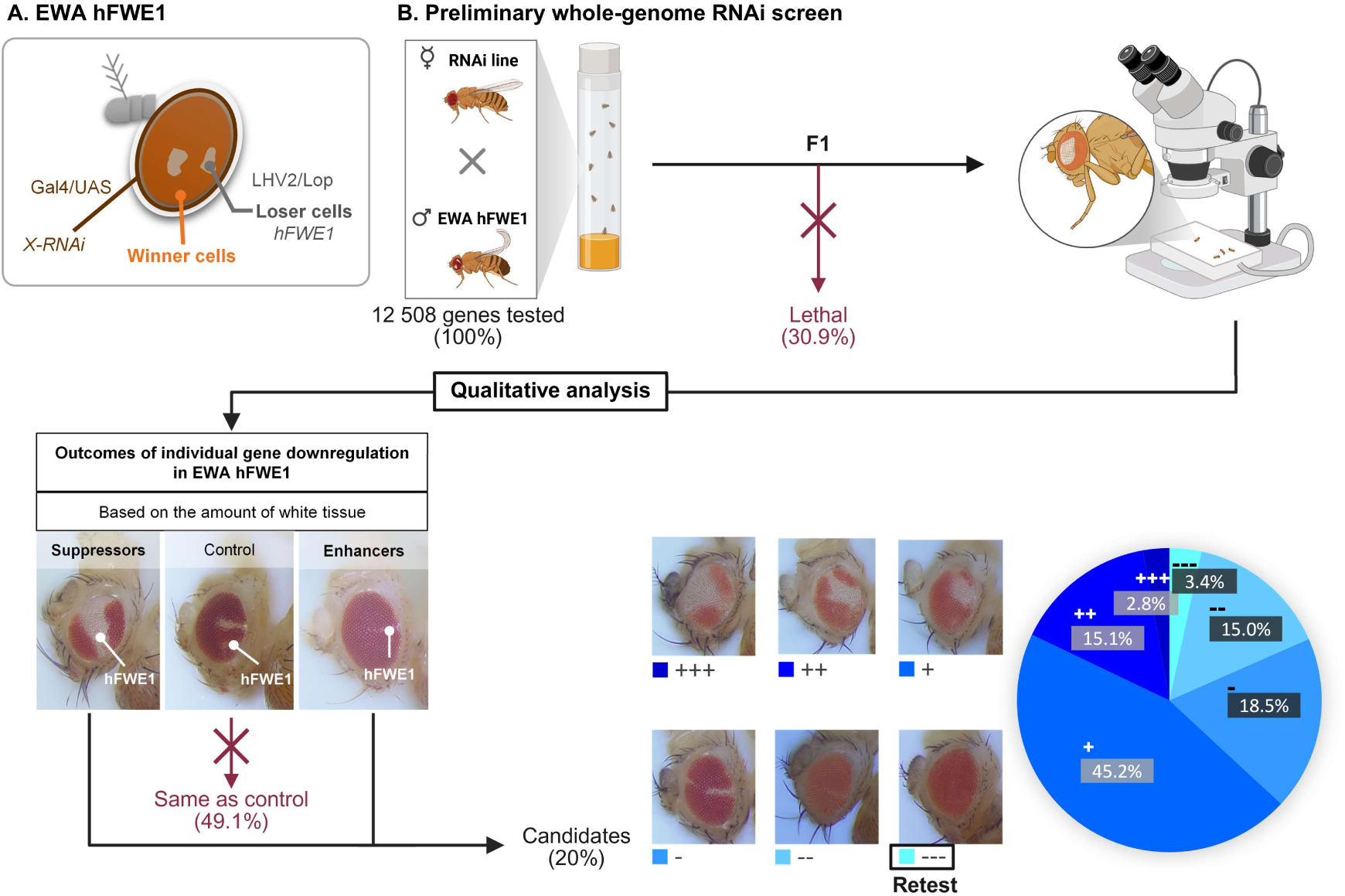
Whole-genome RNAi screen to identify genes required in Fwe-mediated cell competition. **A.** The *LHV2/Lop* clonal subset is labeled in white and corresponds to the *hFWE1*-overexpressing loser cells. *Gal4/UAS* system is active in the entire eye, allowing downregulation of genes (*X RNAi*) simultaneously in *LHV2*-negative (winners) and *LHV2*-positive (losers) cells. **B.** Workflow of the preliminary whole-genome RNAi screen. A total of 12 508 genes (one RNAi line *per* gene) were screened for their requirement in hFWE1-dependent cell competition. 30.9% of lines tested were lethal in this setting. Phenotypic analysis of different outcomes of individual gene downregulation was performed on the progeny of the initial cross, based on the amount of white tissue in the adult eyes. 49.1% of genes tested were classified as being the same as the control and were discarded. The remaining 20% were identified as potential candidate genes, sorted into suppressors and enhancers. These were qualitatively categorized according to the strength of the phenotype. The percentage of each subcategory relative to the total amount of candidates is shown in the diagram. All candidates without clones were retested.

Following this approach, 30.9% of RNAi lines tested were lethal, likely due to developmental defects. Based on visual inspection of the progeny under the stereo microscope, the remaining genes yielded an eye phenotype that was scored blind to the gene tested (**Figure 2B**), with the proportion of white tissue used as a proxy for clone survival or elimination. Tested genes were classified as “same as control”, “suppressor” or “enhancer” according to whether the percentage of white tissue was unaltered, increased or reduced, respectively, compared to *hFWE1* control clones. Among all tested genes, 49.1% were comparable to the control, while 12.6% were identified as suppressors and 7.4% as enhancers. Additionally, genes were rated based on other phenotypic features, independently of the amount of white tissue, such as appearance of black (necrotic) tissue in the eye, or eye malformations (size and/or morphology) specific to the hFWE1 competition setting compared to activation of the RNAi alone (**Table S1 and Figure S3A**). Also, suppressor phenotype resulting in dispersed smaller clones, instead of normally large clones, was categorized as rescue of smaller clones (**Table S1 and Figure S3B**). Lastly, all potential enhancer RNAi lines, in which no white clones were recovered, were retested to minimize false positives that may not show clones due to early activation of *ey-FLP* in the germline. Overall, in the initial screen, we identified 20% of genes tested as candidate modulators of hFWE1 clonal interactions. Suppressors were further categorized from very strong (+++) to mild (+) and enhancers from very strong (---) to mild (-) (**Figure 2B and Table S1**). Among the identified suppressors, we successfully recovered known components of the apoptotic pathway, such as *Death regulator Nedd2-like caspase* (*Dronc*) and *Death-associated APAF1-related killer* (*Dark*). The recovery of these established regulators of cell elimination (Igaki, Yamamoto-Goto, et al., 2002) provides solid validation of our screening strategy.

To establish robust modifiers (**Figure 3A**), we retested very strong suppressors (+++) and further included membrane-associated strong hits (++), given their increased likelihood to interact with Fwe at the membrane, using increased sample sizes and multiple RNAi lines *per* gene whenever possible. Among the 133 retested suppressors, and following quantification (**Figure 3A**), 30 suppressors were retained as high-confidence suppressors of hFWE1-mediated cell elimination (**Figure 3B and Table S2**). Among the 30 suppressors, we retrieved a known binding partner of Fwe, a membrane-anchored heparan sulfate proteoglycan encoded by *division abnormally delayed* (*dally*) (Merino et al., 2022). We further identified genes encoding for the Tumor Necrosis Factor (TNF)-α receptor *grindelwald* (*grnd*) and the scaffolding protein encoded by *veli*, which were previously described to physically interact with each other (Andersen et al., 2015). Among the remaining candidates, we highlight the presence of several metabolism-related genes, including enzymes involved in β-oxidation ( *β Hydroxy acid dehydrogenase 1, Had1*), triglyceride biosynthesis (*CG33120*), chitin biosynthesis (*Chitin synthase 2, Chs2*), and sphingomyelin catabolism (*neutral sphingomyelinase, nSMase*). We also find sodium/solute symporters (*Sodium/solute co-transporter-like 5A11, SLC5A11*, and *CG31262*), potassium channels (*Shaker cognate w*, *Shaw,* and *Shaker, Sh*), and ionotropic receptors (*Ionotropic receptor 76a, Ir76a*, and *Ionotropic receptor 94a, Ir94a*), which may reveal a role for nutrients and ions as modulators of intercellular signaling. Clonal interactions also appear to be regulated by ECM-associated factors (*CG31999*) and regulators of cytoskeleton and adherens junctions (*FER tyrosine kinase*, *FER*). Taken together, these results point to metabolism and intercellular signaling as important modulators of Fwe-driven cell competition.

**Fig. 3.**
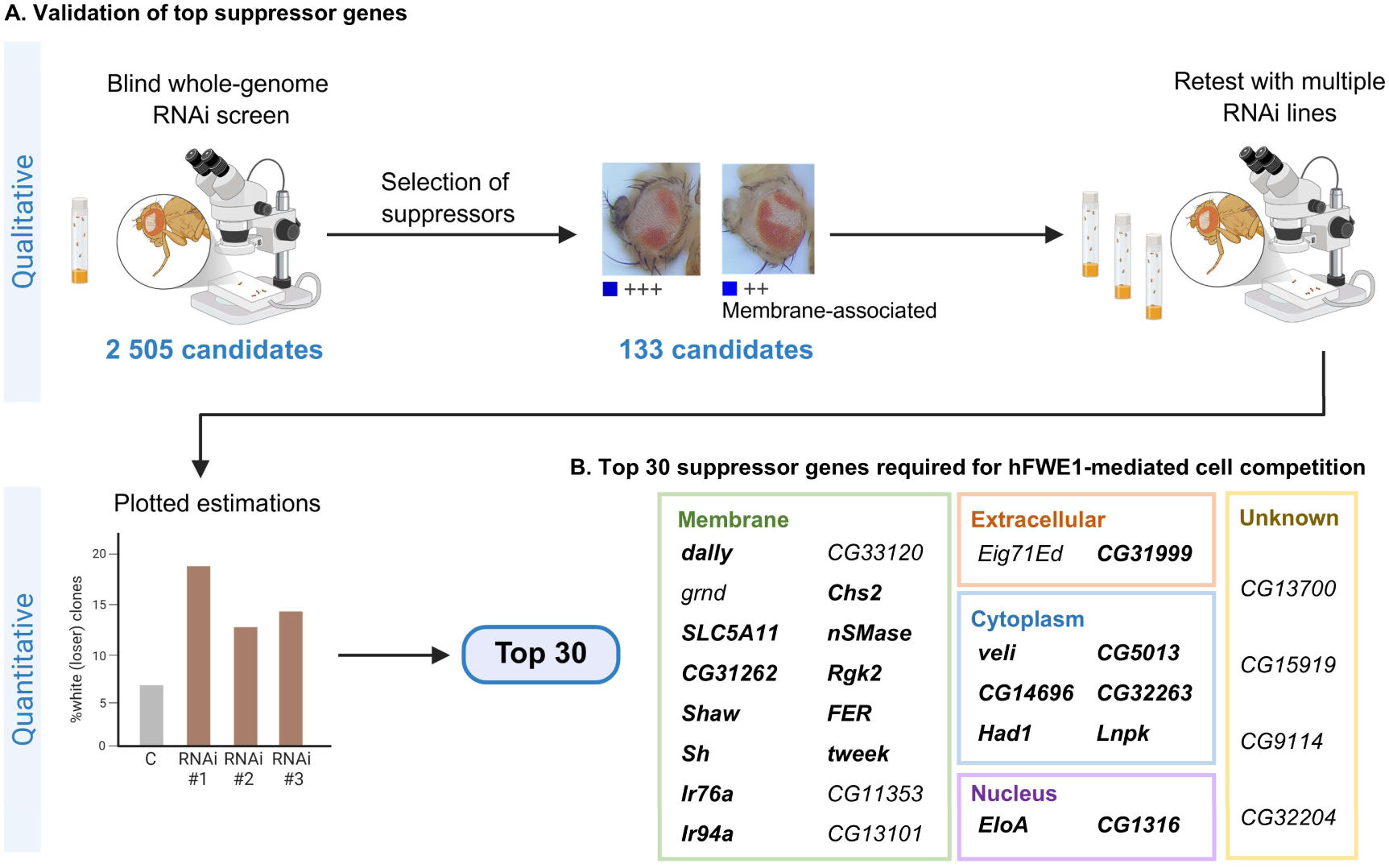
Identification of 30 suppressors of hFWE1-mediated cell competition. **A.** Workflow for the validation of top suppressor candidates. The initial blind whole-genome RNAi screen identified 2 505 candidate genes potentially required for hFWE1-mediated cell competition. Among the 133 suppressor genes selected for further testing, 30 showed statistically significant differences from the control. **B.** The top 30 suppressors required for hFWE1-mediated cell competition are shown, divided by their predicted cellular location. Genes with human orthologs are highlighted in bold.

### *veli* and *grnd* are required for hFWE1-mediated cell competition

Given the joint recovery of the receptor-scaffold Grnd-Veli pair in our screen, we next interrogated how these components act in hFWE1-driven cell competition. Analysis of clonal coverage revealed that downregulation of either *veli* or *grnd* in the entire eye significantly rescued the elimination of *hFWE1* loser cells (**Figure 4A-D**). Furthermore, downregulation of *veli* or *grnd* (*ey>act>Gal4*) alone, in the absence of *hFWE1* overexpression, resulted in an eye of normal size (bottom panels in **Figure 4A,C**). Thus, the observed rescue of *hFWE1* loser clones is not due to an intrinsic growth advantage of *grnd RNAi* or *veli RNAi* tissue. Additionally, to exclude that *veli* or *grnd* act as general regulators of apoptosis, we analyzed cell death-dependent terminalia and spermiduct rotation in male flies (Macías et al., 2004), which was neither affected by *veli* or *grnd RNAi,* nor knockdown of *fwe*. In contrast, overexpression of the cell death inhibitor *Diap1* yielded an inverted, non-functional terminalia as expected (**Figure 4E,F**). Together, these results show that *veli* and *grnd* are required for hFWE1-dependent loser clone elimination, but dispensable for developmentally regulated apoptosis and are, therefore, not generally required for cell death.

**Fig. 4.**
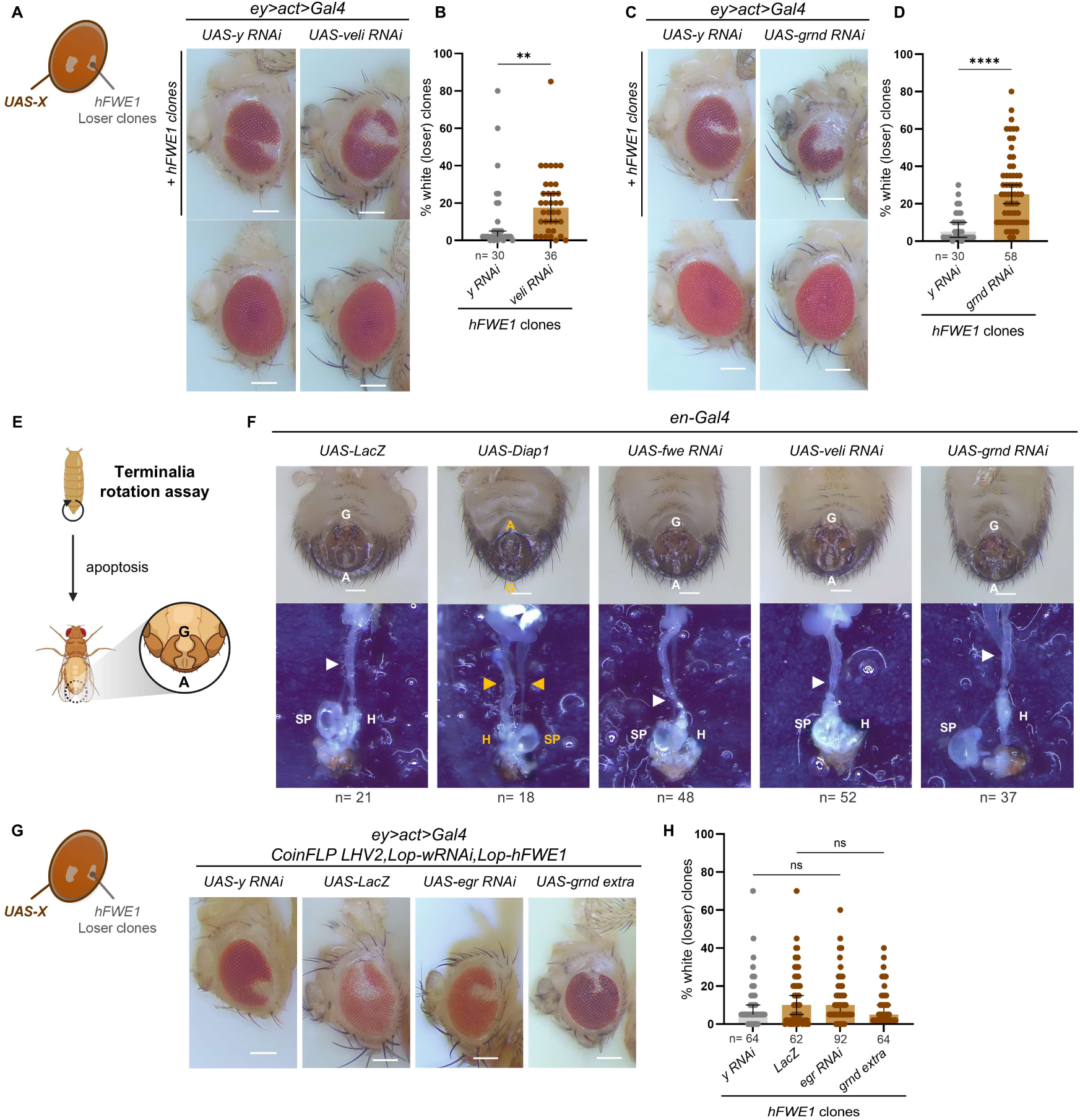
veli and grnd are required for hFWE1-mediated cell competition in a Egr-independent manner. **A,C.** Representative images of downregulation of *veli* or *grnd* in both winner and *hFWE1* loser cells (*ey>CoinFLP LHV2,Lop-hFWE1,Lop-wRNAi; >act>Gal4,UAS-X RNAi*; top panel) and downregulation in the entire eye without clones (*ey>act>Gal4,UAS-X RNAi*; bottom panel). **B,D.** Quantification of the percentage of recovered white loser tissue upon downregulation of *veli* or *grnd* in the entire eye. Crosses at 29°C; n=number of eyes (females); Mann-Whitney test. **E.** Male terminalia rotation assay for cell death. During the pupae stage, cell death is required for proper rotation. Wild-type terminalia is represented (A – analia, G – genitalia). **F.** *Engrailed* (*en*) drives the expression of different transgenes in the terminalia (top panel). Dissected adult males are shown on the bottom panel (SP – sperm pump, H – hindgut), highlighting the spermiduct rotating around the hindgut with a white arrowhead. Absence of this rotation is identified with a yellow arrowhead. Experiment performed at 25°C; n=number of males. **G.** Representative images of adult eyes containing *hFWE1* white loser clones and overexpression of different transgenes in the entire eye (*ey>CoinFLP LHV2,Lop-hFWE1,Lop-wRNAi; >act>Gal4,UAS-X*). **H.** Quantification of the percentage of recovered white loser tissue in each condition. Crosses at 29°C; n=number of eyes (females); Kruskal-Wallis test (multiple comparisons).

### *veli* and *grnd* are required in both winner and loser cells to drive Eiger-independent elimination of hFWE1 loser clones

Next, we interrogated whether *hFWE1* clone elimination is dependent on *Drosophila* TNF-α/Eiger (Egr), the ligand for Grnd (Andersen et al., 2015). Downregulation of Egr function, either by knocking down its expression (*UAS-egr RNAi*), or by overexpressing a dominant negative form of its receptor Grnd that sequesters Egr (*UAS-grnd^extra^*, lacking the intracellular domain), in the entire eye, failed to rescue *hFWE1* clones (**Figure 4G,H**). Therefore, although Grnd is necessary for the elimination of these cells, the receptor functions independently of its canonical ligand Egr in this context. Moreover, when we tested *grnd* and *veli RNAi* in a model of Egr-JNK driven cell death in the eye (Igaki, Kanda, et al., 2002; Moreno et al., 2002), *grnd* suppression yielded a substantial rescue of the triggered small eye phenotype, as well as overexpression of *Diap1*, whereas *veli* inhibition did not show any effect (**Figure S4A-C**), demonstrating that *veli* suppression does not disrupt signaling in the canonical Egr-Grnd-JNK axis in this context.

To further dissect the roles of *grnd* and *veli*, we tested whether these genes are specifically required in loser or winner cells. We first asked whether knocking down their expression in wild-type (WT) clones was sufficient to trigger an effect *per se*. Downregulation of either gene in the absence of cell competition did neither alter clonal size nor eye morphology (**Figure 5A,B**). For exclusive suppression of *veli* and *grnd* in winner cells, we established a modified EWA stock, in which *Lop-Gal80* prevents Gal4 activity in loser *hFWE1*-expressing clones (**Figure S1C**). When RNAi activity was limited to winner tissue (*ey>Gal4;CoinFLP LHV2>Lop-Gal80,Lop-hFWE1*), only *veli* but not *grnd* knockdown significantly rescued *hFWE1* loser clones, revealing that *veli* is required in winners to drive Fwe-dependent cell elimination of neighboring loser cells (**Figure 5C,D**). Next, to manipulate *grnd* and *veli* function specifically in loser cells, we used a Gal4-dependent CoinFLP system. We confirmed that inhibiting apoptosis, either by expression of *Dronc RNAi* (**Figure 5E,F**) or overexpression of *Diap1* (**Figure 5G,H**), successfully prevents cell elimination. In addition, exclusive suppression of *veli* in loser clones prevented their elimination. Intriguingly, knockdown of *grnd* in losers failed to rescue them (**Figure 5E,F**). Additionally, overexpression of the dominant negative *grnd^extra^* in loser cells did not prevent their elimination (**Figure 5G,H**), further supporting our previous conclusion that loser cell death is Egr-independent. Taken together, these results show that Veli can regulate loser elimination in a non-cell-autonomous manner, with its function required in both loser and winner cells to drive Fwe-dependent cell competition. In contrast, Grnd plays a non-canonical, Egr-independent role. Loss of *grnd* in either cell population can be compensated for by the other, whereas a tissue-wide depletion of *grnd* prevents hFWE1-dependent cell elimination altogether.

**Fig. 5.**
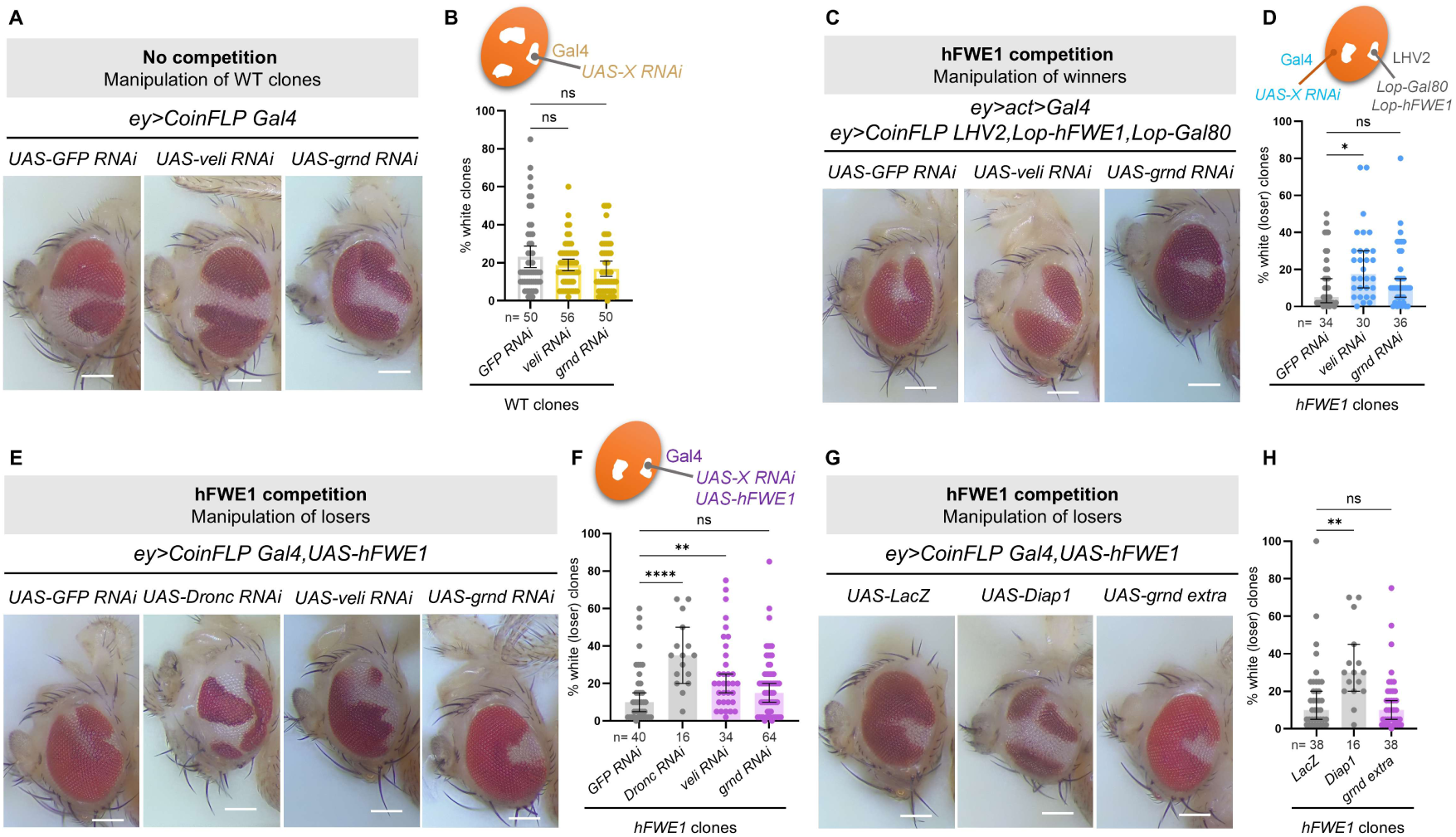
veli and grnd are required in both winner and loser cells for the elimination of hFWE1 losers, but in distinct manners. **A**. Downregulation of control *GFP*, *veli* or *grnd* in non-competitive WT clones (*ey>CoinFLP Gal4,UAS-X RNAi, UAS-wRNAi*). **B.** Quantification of percentage of white clones. **C.** Downregulation of control *GFP*, *veli* or *grnd* in the winner population (*ey>CoinFLP LHV2,Lop-hFWE1,Lop-Gal80,Lop-wRNAi; >act>Gal4,UAS-X RNAi*). **D.** Quantification of the percentage of the recovered amount of loser tissue upon downregulation of transgenes in the winner population. **E.** Downregulation of control *GFP*, *veli* or *grnd* in *hFWE1* loser clones (*ey>CoinFLP Gal4,UAS-hFWE1,UAS-X, UAS-wRNAi*). **F.** Quantification of percentage of recovered white loser clones. Crosses at 29°C; n=number of eyes (females); Kruskal-Wallis test (multiple comparisons). **G.** Overexpression of control *LacZ*, *Diap1* or *grnd extra* in *hFWE1* loser clones (*ey>CoinFLP Gal4,UAS-hFWE1,UAS-X, UAS-wRNAi*). **H.** Quantification of percentage of recovered white loser clones. Crosses at 29°C; n=number of eyes (females); Kruskal-Wallis test (multiple comparisons).

### Fwe-Grnd-Veli communication module at the winner-loser interface conveys fitness information

To better understand how Grnd and Veli implement their bilateral roles during competition, we generated clones in developing wing discs and monitored the subcellular localization of these proteins. GFP-labeled clones were induced by heat-shock and wing discs were imaged 48 h and 72 h after clone induction (*hsFLP;CoinFLP Gal4,UAS-GFP;UAS-X*) (**Figure 6A**). Under homeostatic conditions, both Grnd and Veli predominantly localize apically, as previously described (Andersen et al., 2015; Bachmann et al., 2004), whereas FweUbi is distributed along the entire cell membrane (**Figure 6B,C and Figure S5A,B**). When hFWE1-dependent cell competition is induced, we detected basal Grnd-positive puncta in loser clones (**Figure 6D,E**), likely representing vesicles preceding cell elimination (Palmerini et al., 2021), alongside apical enrichment of Grnd (**Figure 6D,E and Figure S5C-F**). Strikingly, we also found a strong apico-lateral accumulation of Grnd at winner-loser interfaces, which has not been previously described (**Figure 6D,E**). Similarly, we found enrichment of Veli and FweUbi at the lateral interface (**Figure 6D,E and Figure S5C-F**).

**Fig. 6.**
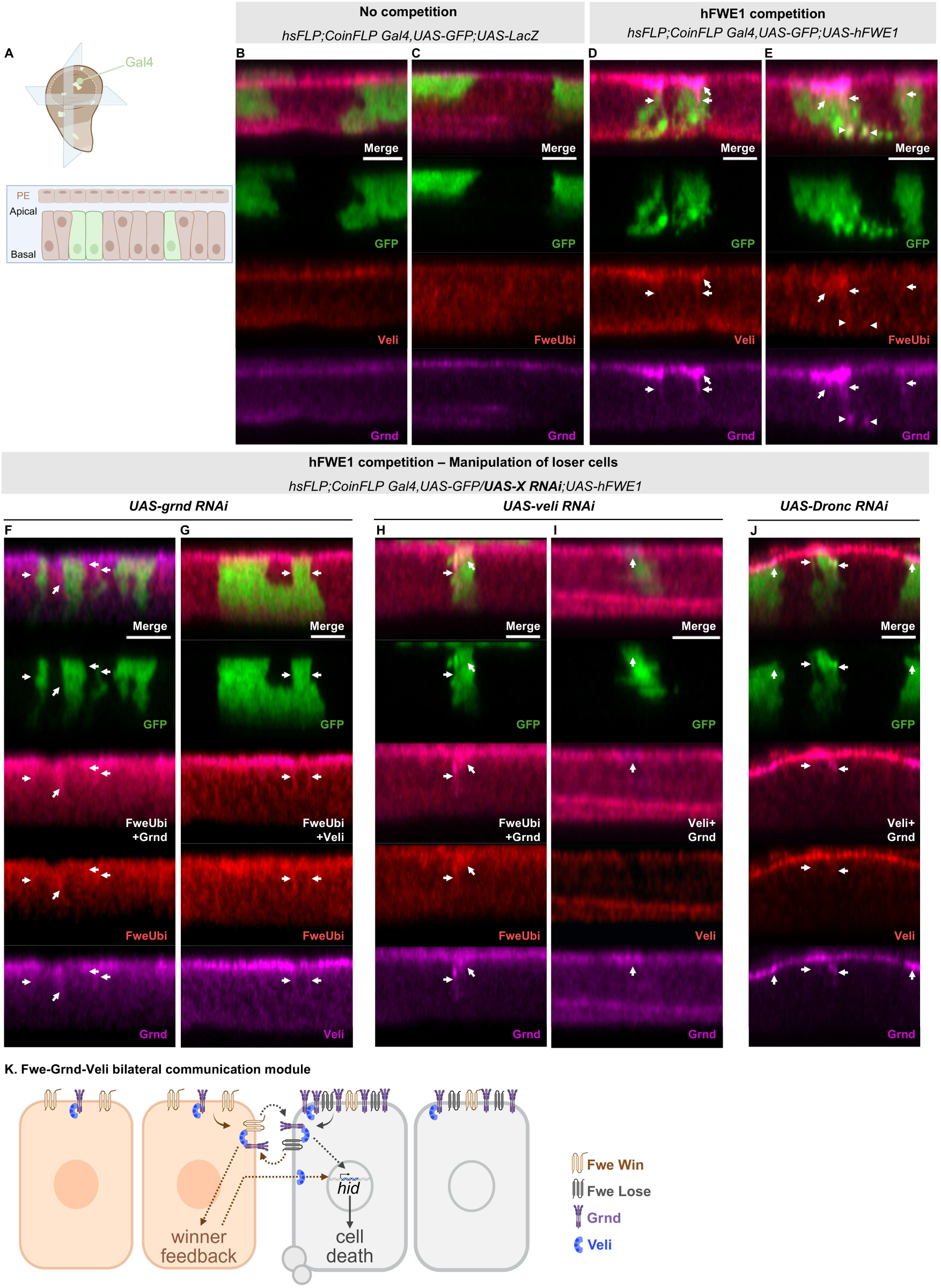
Fwe-Grnd-Veli form a bilateral module at the winner-loser interface. **A.** Wing imaginal discs with GFP-labeled clones (*hsFLP;CoinFLP Gal4,UAS-GFP;UAS-X*). XZ/YZ cross-sections are shown, depicting the pseudostratified epithelium of the disc, with the peripodial epithelium (PE) on top. **B-C.** Veli, Grnd and FweUbi signal in the absence of cell competition (*UAS-LacZ* clones). **D-E.** Veli, Grnd and FweUbi signal in hFWE1-induced competition. Grnd-positive basal puncta are indicated by arrowheads and protein enrichment is indicated by arrows. **F-G.** Veli, Grnd and FweUbi signal in hFWE1-induced competition with downregulation of *grnd* in loser clones. Protein enrichment is indicated by arrows. **H-I.** Veli and Grnd signal in hFWE1-induced competition with downregulation of *Dronc* in loser clones. Protein enrichment is indicated by arrows. **J.** Veli, Grnd and FweUbi signal in hFWE1-induced competition with downregulation of *grnd* in loser clones. Protein enrichment is indicated by arrows. Cross-sections at 20x magnification of the pouch of wing imaginal discs are shown, 48 h and 72 h after clone induction. n>10 wing discs were analyzed for each condition. FweUbi, Grnd and Veli were detected with antibodies and are color-labeled as indicated in each panel. **K.** Working model for Fwe-Grnd-Veli bilateral communication module. When Fwe competition is induced and the communication module is correctly assembled, Grnd becomes apically enriched in loser cells and redistributes to the winner-loser interface in both winners and losers. Loser-side and winner-side Grnd sense neighbors’ fitness. In loser cells, Veli integrates Fwe Lose and Grnd signal and activates the loser response, culminating in downstream upregulation of the pro-apoptotic gene *hid* and cell death. At the same time, Veli in winners integrates FweWin and Grnd signal and activates the winner feedback program, which subsequently enhances elimination of the neighboring loser cell. On the other hand, when the communication module lacks Veli on either side, it suppresses the integration of relative fitness differences. Simultaneous depletion of Grnd in loser and winner cells does not allow neighboring fitness sensing. Both cases impede activation of loser cell death.

Next, we downregulated *veli* or *grnd* in loser cells to examine effects on the observed protein redistribution in the distinct membrane domains. Downregulation of *grnd* expectedly reduced Grnd signal in loser clones (**Figure 6F and Figure S5G**), but Grnd, Veli and FweUbi could still be jointly detected at the winner-loser interface, indicating that winners can considerably contribute to Grnd localization at the clonal interface (**Figure 6F,G and Figure S5G,H**). When *veli* is downregulated in loser clones, resulting in Veli depletion (**Figure 6I and Figure S5J**), Grnd still remains apically enriched in loser cells and can be detected in the winner-loser interface (**Figure 6H,I and Figure S5I,J**). This shows that Veli is not required for Grnd localization or redistribution, in line with previous findings (Andersen et al., 2015), and that the accumulation and redistribution of Grnd is not sufficient to trigger elimination when *veli* is absent. Interestingly, loser cells with *veli* knockdown show an accumulation of FweUbi, normally associated with increased fitness and “winner” outcome (**Figure 6H**), suggesting altered cellular interpretation of fitness state, in line with the observed reduced elimination of these clones. Lastly, we asked whether blocking apoptosis of loser cells would impact the protein recruitment to the winner-loser interface. Knockdown of *Dronc* did not prevent Grnd clustering apically and at interfaces, with some Veli accumulation also being detected at the interface (**Figure 6J and Figure S5K**), similar to *hFWE1*-control clones, indicating that this enrichment occurs upstream of caspase activation. Overall, these findings suggest a model in which Fwe, Grnd and Veli are dynamically redistributed during cell competition and assemble into a bilateral interface module, enabling bidirectional fitness comparison at the winner-loser interface to promote the recognition and elimination of loser cells (**Figure 6K**).

## Discussion

Cell competition relies on the ability of neighboring cells to compare relative fitness and eliminate the ones with lower fitness for the benefit of optimal organ function, for which Flower (Fwe) fitness fingerprints have been shown to play an important role (Madan et al., 2019; Rhiner et al., 2010). In this study, we developed a customizable clonal interaction assay in *Drosophila*, the Easy Win Assay (EWA), to conduct a genome-wide RNAi screen for novel modulators of hFWE1-induced cell elimination. Among the candidate genes successfully validated, we observed a strong enrichment of factors involved in metabolism and signaling. In particular, we uncover that the receptor Grindelwald (Grnd) and the scaffold protein Veli are central components of the winner-loser interface mediating fitness comparison, within which they act as bilateral mediators of hFWE1-dependent cell elimination. As key features of this module, we establish a bilateral requirement of Veli in loser and winner cells, plus interface presence of Grnd, which can be contributed by either winner or loser cells to allow removal of hFWE1-flagged losers. These findings underscore that winner and loser cells do not simply play asymmetric roles as “killer” and “victim”, but instead jointly contribute to the formation of a signaling module that integrates and coordinates fitness comparison.

Grnd, one of the *Drosophila* TNF receptors, has been shown to bind Eiger (Egr)/TNF-α and act upstream of JNK-dependent activation of apoptosis (Andersen et al., 2015), including in competition contexts such as the elimination of NR2-depleted losers (Banreti & Meier, 2020) and polarity-deficient *scrib^-/-^* clones (Andersen et al., 2015; Igaki et al., 2009). On the other hand, wild-type cells outcompeted by Myc-overexpressing cells are largely eliminated independent of JNK signaling, but require Egr-Grnd (Kodra et al., 2024). Our findings provide strong evidence that Grnd operates non-canonically during Fwe-dependent cell competition. Rather than acting as a simple loser-autonomous death receptor, Grnd emerges as an essential, Egr-independent component of Fwe-driven cell elimination that must be functional in at least one of the apposed clonal populations with differential fitness. We propose that Grnd functions as a fitness sensor that integrates Fwe-dependent signals at the winner-loser interface. Our data further suggest that this integration occurs at specialized lateral membrane domains, enabling direct comparison of winner and loser cell membrane states. During competition, Grnd, Veli and Fwe are laterally redistributed to the winner-loser interface, supporting a model in which these components assemble into a dynamic signaling module that mediates fitness comparison and activates downstream responses. We envisage that the localization of this module to the lateral membrane domain represents a crucial step in driving the fitness pathway activation. It remains to be determined whether this module acts directly in the initial fitness recognition step or is engaged downstream once differential fitness is recognized.

Here, we identify a previously unappreciated role for Veli in epithelial quality control, where it is required in both winner and loser cells for Fwe-dependent competition to be active. Notably, although loss of *veli* does not impair apico-lateral recruitment of Grnd, it leads to increased FweUbi levels in loser cells, suggesting that Veli is not required for the assembly of the module but instead integrates information and modulates downstream signaling. This view is consistent with Veli’s role as a scaffolding protein, which while not required to establish or maintain cell polarity, couples apical complexes to downstream processes in epithelia and neurons (Bachmann et al., 2004, 2008; Soukup et al., 2013). Through its established physical interaction with Grnd, it can relay membrane-associated changes to downstream intercellular signaling pathways, similar to polarity-deficient contexts, where Grnd couples Crumbs-dependent apico-basal polarity to JNK signaling and neoplastic growth (Andersen et al., 2015). Together with its beneficial function in postsynaptic neurons to prevent light-induced photoreceptor degeneration (Soukup et al., 2013), these observations support a model in which Veli acts as a context-dependent signaling hub that coordinates pruning or resilience under stress-induced conditions.

Other potential downstream responses or parallel pathways impinging on Fwe-dependent cell competition could involve metabolism, ion transport and extracellular matrix (ECM) organization, as hinted by the validated suppressors from the screen. This is consistent with the emerging view that metabolic state, microenvironment and membrane-associated components influence competitive outcomes (Banreti & Meier, 2020; Esteban-Martínez & Torres, 2021; Morata, 2021; Nagata & Igaki, 2024). Additional phenotypes identified in the screen, such as the formation of necrotic or melanotic tissue, may reflect impaired tumor suppression, whereas suppressors that promoted survival of small, scattered clones point towards robust pathway blockage or altered cell mixing properties (Levayer et al., 2015). Lastly, because many of these regulators are conserved in mammals, our dataset also provides a valuable foundation for investigating these mediators in mammalian co-culture or organoid models of cell competition.

Beyond genetic screens, the versatile EWA system provides a broad platform to explore non-genetic modulators of cell-cell interactions (Costa et al., 2020) such as pharmacological compounds or nutrient perturbations. Furthermore, altering the timing or tissue specificity of recombinase expression enables clonal induction across diverse tissues, establishing a powerful approach to dissect diverse aspects of intercellular communication.

Future studies will be essential to determine whether the modulators of hFWE1-induced competition identified here are broadly required across other cell competition paradigms, such as Xrp1-mediated, polarity-defective or Myc-driven competition. Resolving this will provide further insight into the long-standing question in the field of whether different forms of cell competition rely on common or distinct mechanisms (Lavalou et al., 2025; Matsumoto et al., 2026; Morata, 2021).

Once the decision is made to eliminate a suboptimal cell, our results suggest that the execution machinery can be shared across different cell elimination processes. Fwe-dependent cell competition and tumor-suppressive competition share a downstream module, yet they are triggered by different fitness-sensing inputs. These upstream recognition mechanisms can independently regulate cell elimination, giving the organism flexibility in the identification and removal of diverse damaged or dangerous cells. Fwe-dependent cell elimination is targeted towards preventing aging-related decline and developmental malformations but can promote tumor growth through the appearance of supercompetitor cells, rather than acting as a tumor-suppressive pathway (Costa-Rodrigues et al., 2021; Gutiérrez-Martínez et al., 2019; Vishwakarma & Piddini, 2020). This natural distinction may have therapeutic implications. For example, modulating upstream regulators of cell selection like Fwe, rather than more common apoptotic pathways, could offer a cleaner way to regulate aging, development and tumor invasion, while potentially not affecting tumor-suppressive death pathways.

Overall, our findings provide mechanistic insight into how neighboring epithelial cells compare relative fitness and eliminate suboptimal cells during Fwe-dependent cell competition. We demonstrate that both winner and loser cells actively drive this process, with a newly identified interface signaling module, comprising Fwe, Grnd and Veli, serving as a central determinant of cellular competition. This work advances our understanding of tissue-level quality control and provides a framework to explore new avenues for modulating competitive interactions in disease contexts where fitness levels are altered.

## Limitations of the study

Although allowing for easy detection of clonal outcome in the eye tissue, the early induction of clones driven by the *eyeless* promoter can interfere with early development and obstruct phenotypes, as evidenced by the considerable number of RNAi lines resulting in lethality. The human FWE1 Lose isoform was selected for its highly robust effect in triggering cell elimination. However, it remains to be determined how *Drosophila* cells integrate the overexpression of this human isoform.

## Acknowledgments

We thank Sandra Soukup, Julien Colombani and Ditte Andersen for antibodies, Konrad Basler for plasmid, Manuel Calleja, Hugo Stocker, Julien Colombani, Ditte Andersen, Pierre Léopold, Norbert Perrimon, the Vienna Drosophila Resource Center (VDRC) and the Bloomington Drosophila Stock Center (BDSC) for fly lines, and the Champalimaud Foundation Molecular Tools and Transgenics Platform and Imaging (ABBE) platform for technical support. Figure schemes were created in https://BioRender.com.

## Author contributions

Conceptualization: EM. Methodology and Investigation: CBP established and validated the EWA-method, CMC, JC, DC, MCA conducted the screen, SB performed apoptosis assays, and CMC performed the remaining experiments. Validation, formal analysis and visualization: CMC. Writing: CMC with contributions from CR, CBP and EM. Supervision: CR, EM. Funding acquisition: EM, CBP and CMC.

## Supplemental information

Document S1. Figures S1-S5.

Table S1. Results from the initial whole-genome RNAi screen.

Table S2. Quantification analysis of 30 validated suppressors of hFWE1-induced cell competition.

## Materials and Methods

### Fly strains and husbandry

Flies (*Drosophila melanogaster*) were raised on yeast-based Vienna food, under controlled temperature (22°C or 25°C) and humidity (70% relative humidity), on a 12h light cycle. Experimental crosses were set up at 25°C or 29°C as indicated in each figure. The following fly stocks were used: *yw,eyFLP,act>STOP>Gal4* (kind gift from Hugo Stocker,(Nowak et al., 2018)), *eyFLP,UAS-Dcr2* (BDSC#58756), *CoinFLP Gal4,UAS-GFP* (BDSC#58751), *CoinFLP Gal4* (BDSC#58750), *CoinFLP LHV2* (this work), *en-Gal4* (BDSC#1973), *tub-Gal4*, *GMR-Gal4*, Lop*-wRNAi* (this work), *Lop-GFP* (BDSC#32207), *Lop-Gal80* (BDSC#32215), *Lop-Myc* (this work), *Lop-hid* (kind gift from Manuel Calleja), *Lop-scrib RNAi* (this work), *Lop-hFWE1* (this work), *UAS-y RNAi* (VDRC#106068), *UAS-SPARC RNAi* (VDRC#100566), *UAS-fwe RNAi* (VDRC#39596), *UAS-veli RNAi* (VDRC#110556), *UAS-grnd RNAi* (VDRC#104538), *UAS-egr RNAi* (VDRC#45253), *UAS-GFP RNAi* (VDRC#60103), *UAS-Dronc RNAi* (VDRC#100424), *UAS-mCherry*, *UAS-Diap1* (BDSC#6657), *UAS-LacZ*, *UAS-grnd extra* (kind gift from Julien Colombani and Ditte Andersen, (Andersen et al., 2015)), *UAS-CD8:GFP*, *UAS^40D^* (VDRC#60101), *UAS-hFWE1* (Costa-Rodrigues et al., 2025). Additionally, for the whole-genome RNAi screen, *UAS-RNAi* lines were ordered from VDRC (line identifiers are listed in Table S1) (Dietzl et al., 2007).

### Generation of transgenic flies

To generate the CoinFLP-LHV2 plasmid, *CoinFLP-AgeI* backbone plasmid was first generated to introduce a unique *AgeI* restriction site by replacing the *EcoRI-GalL4-attP-SapI* fragment of *CoinFLP-Gal4* (Addgene#52889, (Bosch et al., 2015)) with an *EcoRI-AgeI-attP-SapI* sequence. Subsequently, an *EcoRI-LHV2-AgeI* insert was cloned into the *CoinFLP-AgeI* vector. The LHV2 sequence was amplified from *Dpp-attP-LHV2* plasmid (Yagi et al., 2010), kindly provided by Konrad Basler. Hairpin were generated by annealing complementary oligos containing *XhoI* and *XbaI* restriction sites. The annealed products were cloned into the *26xLexAop2-mCD8::GFP* vector, which had been digested with *XhoI* and *XbaI* to excise the *mCD8::GFP*. Specifically, the SH00017.N and SH02077.N hairpins were inserted to generate *26xLexAop2-wRNAi* and *26xLexAop2-scribRNAi*, respectively. To construct *26xLexAop2-Myc*, the coding sequence was amplified from BDGP clone LD32539. Similarly, the *hFWE1* sequence was amplified from vectors previously described (Madan et al., 2019) and cloned using the same strategy to generate *26xLexAop2-hFWE1* vector.

To generate transgenic flies, the following attP landing-site stocks were used: *attP40* (a kind gift from Norbert Perrimon), *attP2* (BDSC#8622), *attPZH-51C* (BDSC#24482) and *attPZH-86Fb* (BDSC#24749). Plasmids were microinjected into specific landing sites as follows: *CoinFLP-LHV2*, *26xLexAop2-wRNAi*, and *26xLexAop2-hFWE1* were injected into *attPZH-51C*, *attP40*, and *attP2*, respectively. *26xLexAop2-Myc* and *26xLexAop2-scribRNAi* were injected into *attPZH-86Fb*. The *attP*-associated selection markers, namely *y+* (for *attP40* and a*ttP2*) or *3xP3-RFP* (*attPZH-51C* and *attPZH-86Fb*), were removed by crossing the transgenic lines with a Cre recombinase stock (BDSC#34516).

### Whole-genome RNAi screen

One RNAi line *per* gene was tested, as indicated in Table S1. The VDRC KK library was the main RNAi collection used, with the GD and shRNA libraries as alternatives when a KK line was not available. Experimental crosses were kept at 29°C and were performed using RNAi virgins and EWA hFWE1 males, except when the RNAi was associated with the Y chromosome. In the latter case, male progeny was scored against male control, instead of females. The initial screen was unbiased to the gene tested. Subsequent validation steps were also unbiased until after the quantification analysis was performed. For the validation of suppressor candidate genes, a subset list of “membrane-associated candidates” was retrieved from FlyBase (Öztürk-Çolak et al., 2024) using the “batch download” feature, using as input the CG code and output solely the “cellular component”. We used FlyBase (Öztürk-Çolak et al., 2024) and associated websites to find information about the genes.

### Heat-shock clone induction

To optimize vial crowdedness and heat transfer homogeneity, fly crosses were set up at 25°C in narrow vials, with 6-9 virgins and a 3:1 virgin/male ratio. 4-5 days after egg laying, clones were induced in larvae by placing the vials in a water bath at 37°C for 8 minutes. Larvae were kept at 25°C following heat-shock. Late/wandering third instar female larvae were collected at defined time points after clone induction and selected for GFP-positive signal in a ZEISS Discovery.V8 fluorescence microscope.

### Dissection, fixation and immunostaining of imaginal tissues

For wing imaginal discs, staining was performed by dissecting selected larvae in phosphate buffer (PBS) on ice, protected from light sources. Samples were then fixed in 3.7% FA in PBS for 20 minutes at room temperature (RT), washed in PBS containing 0.1% Triton X-100 (0.1% PBT) and incubated with primary antibodies in 0.1% PBT supplemented with 10% normal donkey serum (Jackson ImmunoResearch 017-000-121) for 2 hours at RT. Following washes with 0.1% PBT, samples were incubated for 1 hour at RT with secondary antibodies in 0.1% PBT. Samples were then washed with PBS, incubated with DAPI in PBS (1:1000, Sigma D9542) and wing imaginal discs were mounted in vectashield mounting medium (Vector H-1000). The following primary antibodies were used: anti-FweUbi M (1:30, (Rhiner et al., 2010)), anti-Grnd GP (1:100, a kind gift from Julien Colombani and Ditte Andersen, (Andersen et al., 2015)) and anti-Veli/dLin-7 RB (1:1000, a kind gift from Sandra Soukup, (Bachmann et al., 2004)). The following secondary antibodies were used at a 1:1000 dilution: anti-Rb-A647 (Invitrogen A-31573), anti-Rb-A568 (Invitrogen A-10042), anti-M-A568 (Invitrogen A-10037), anti-GP-A647 (Invitrogen A-21450).

### Image acquisition

Fluorescent images of fixed samples were obtained on a Zeiss LSM 880 confocal laser scanning microscope with ZEN 2.3 software, using 20x ZEISS Plan-APOCHROMAT (20x/0.8).

Images of adult flies’ eyes and male terminalia were taken in a Leica Stereozoom S9i with camera (Leica LED3000 DI), with the Z-axis reconstruction software (LAS X software with LAS X Live Image Builder Z). Flies were kept alive and anesthetized with CO_2_ during image acquisition. Images of dissected males were taken immediately after dissection.

## Quantification and Statistics

For estimation of the percentage of white clones in adult eyes, trained observers used a semi-quantitative scoring approach based on direct visual estimation of the white tissue fraction in relation to the total area of the eye. Quantifications of eye area in the GMR>Eiger assay and of clonal area in wing discs were made using ImageJ (Fiji). For wing discs, maximum intensity projections were generated, excluding the peripodial epithelium. GFP area in the wing pouch was quantified and normalized to the total area of the pouch. For generation of intensity plots, ZEN software was used - apical slices of wing discs were chosen, and a line was drawn to include a clone, its borders and neighboring tissue.

GraphPad Prism 11 was used for statistical analysis. All data were first analysed for normal distribution. Subsequently, parametric or non-parametric statistical analysis was performed, as indicated in each figure. Correction for multiple comparisons was applied where indicated. Statistical significance was defined by a P-value≤0.05. ns, non-significant P>0.05, * P≤0.05, ** P≤0.01, *** P≤0.001, **** P≤0.0001. Graphics plot the median with 95% confidence interval for error bars for non-normal distributions, and mean with standard deviation for normal distributions.

## Resource availability

### Lead contact

Requests for further information and resources should be directed to and will be fulfilled by the lead contact, Eduardo Moreno.

### Materials availability

Transgenics generated in this study will be made available upon request to the lead contact.

### Data and code availability

- Data reported in this paper is included as main and supplementary data.
- This paper does not report original code.
- Any additional information required to reanalyse the data reported in this paper is available from the lead contact upon request.

