## Supplementary material for "A whole-genome screen identifies a bilateral Flower-Grindelwald/TNFR-Veli/LIN-7 communication module in cell competition": Document S1

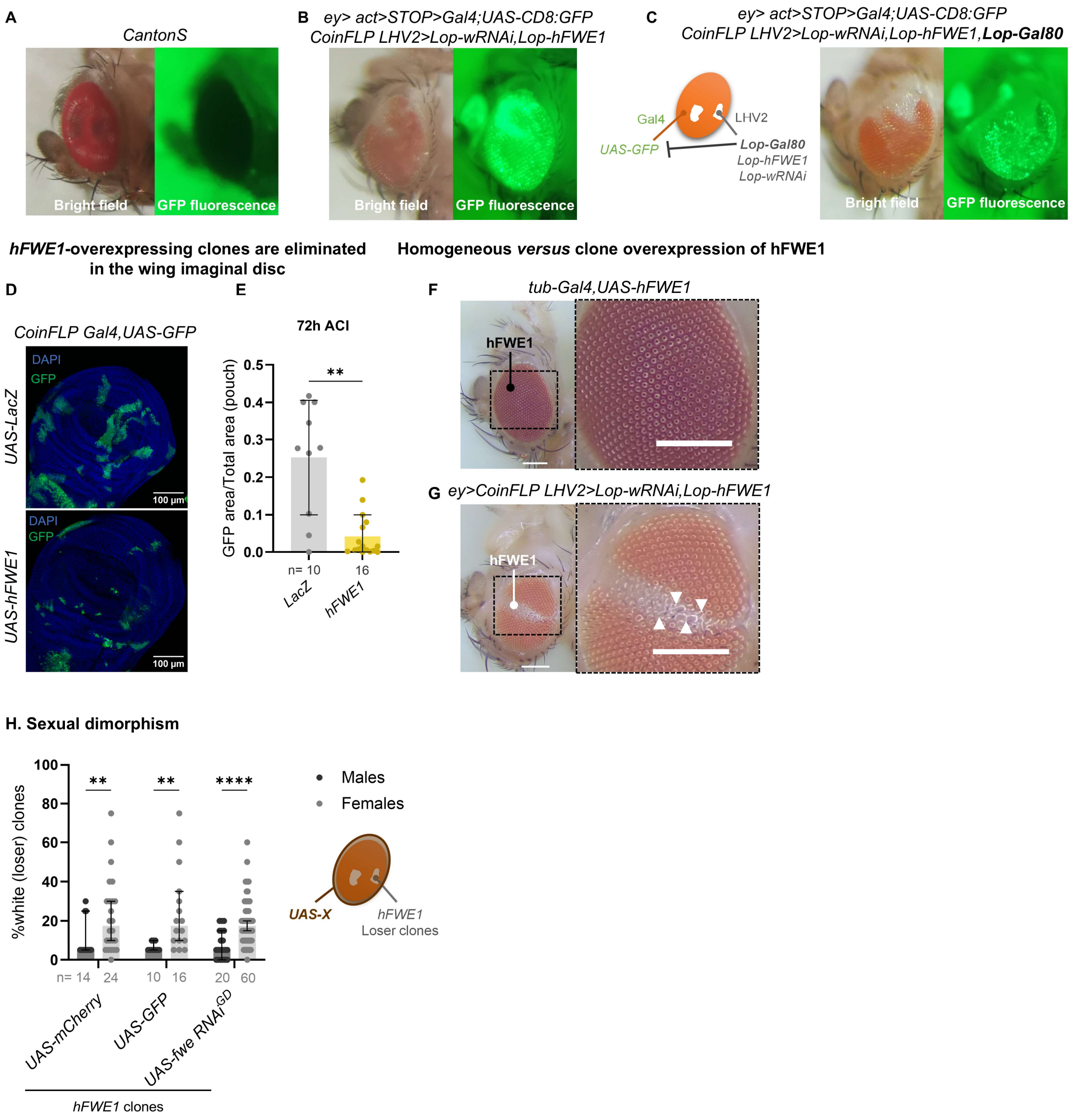


**Supplemental Fig.1 – A-C.** Bright field and GFP fluorescence images of adult flies are shown. **A.** Wild-type (*CantonS*) flies do not have green autofluorescence. **B.** When crossing EWA hFWE1 flies with *UAS-CD8:GFP*, GFP signal can be detected throughout the entire eye and antennae, demonstrating that the *act>STOP>Gal4* cassette flips completely. **C.** In turn, when LHV2 clones express *Lop-Gal80*, it represses Gal4 activity in these clones, which can be confirmed by the lack of GFP signal in LHV2 white-labelled clones. The surrounding tissue maintains Gal4 activity and positive GFP signal. **D.** ***hFWE1*-overexpressing clones are eliminated in the wing imaginal disc.** GFP-labeled *hFWE1*-overexpressing clones are eliminated from the imaginal wing pouch 72h after clone induction (ACI) (*hsFLP>CoinFLP Gal4,UAS-GFP,UAS-X*). Maximum intensity projections are shown at 20x magnification. Nuclei are shown in blue (DAPI) and clones are shown in green. **E.** GFP area in the pouch was measured and normalized to the pouch area.  Crosses at 25°C; n=number of wing discs (females); Unpaired t test. **F-G.**

**Homogeneous *versus* clone overexpression of hFWE1.** Overexpression of *hFWE1* in the entire tissue (*tub*>) produces a normal eye, while overexpression of *hFWE1* in clones (*ey>CoinFLP LHV2*) gives rise to a rough and disorganized eye tissue (exemplified by white arrowheads). Zoomed-in areas are outlined by black dashed boxes. **H. Sexual dimorphism.** Males consistently present a lower percentage of recovered white loser clones compared to females, across different conditions in the EWA hFWE1 (*ey>CoinFLP LHV2,Lop-hFWE1,Lop-wRNAi; >act>Gal4,UAS-X*). Crosses at 25ºC; n=number of eyes; Multiple Mann-Whitney test corrected for multiple comparisons.

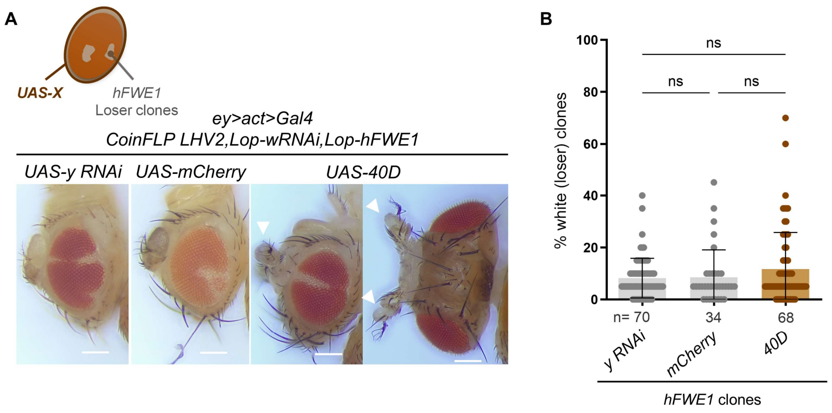


**Supplemental Fig.2 – Control for extra insertion of KK lines.** It is reported that approximately 25% of KK lines from VDRC have an extra insertion at *40D,* which leads to ectopic expression of *tiptop* gene. To dissect the possible effect of this extra insertion on the results of the screen, the tester line for *40D* was compared to other controls used in the EWA (*ey>CoinFLP LHV2,Lop-hFWE1,Lop-wRNAi; >act>Gal4,UAS-X*). Although it does not affect the amount of recovered white tissue and, consequently, the elimination of *hFWE1* loser cells, the extra insertion at *40D* generated a phenotype of big antennas in adult flies (indicated by white arrowheads). Crosses at 29ºC; n=number of eyes (females); Kruskal-Wallis test (multiple comparisons).

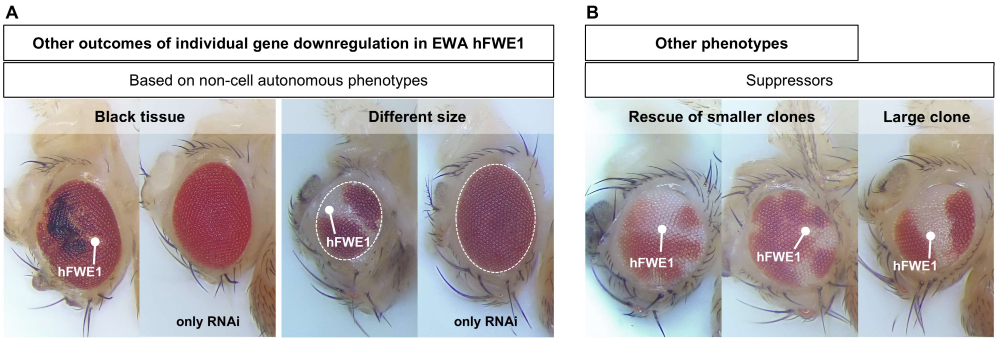


**Supplemental Fig.3 – Other phenotypes observed in the EWA hFWE1. A. Based on non-cell autonomous phenotypes.** In the initial screen, some non-cell autonomous phenotypes were observed upon the downregulation of a gene solely in the EWA hFWE1 setting, and not by the downregulation of the gene alone. These include appearance of black tissue, and malformations regarding size or tissue. **B. Other phenotypes.** Suppressors usually present one large white area, but in some candidates, the recovery of dispersed white clones could be detected, designated here as rescue of smaller clones. In some cases, the white tissue has normal morphology, in contrast to the usually observed rough white loser tissue.

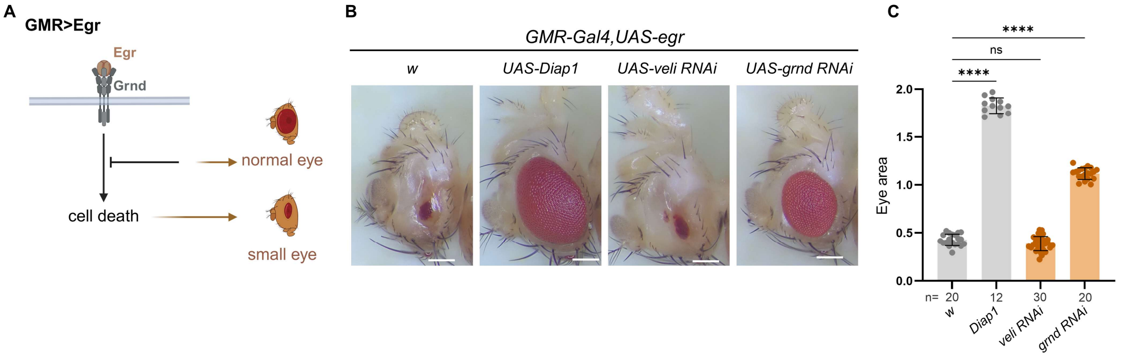


**Supplemental Fig.4 – GMR>Eiger eye assay for cell death. A. Eiger-induced cell death assay.** The ligand Egr binds its receptor Grnd, inducing downstream cell death, producing a small adult eye (*GMR-Gal4,UAS-egr*). If cell death is blocked, the eye size is rescued. **B,C.** Overexpression of *Diap1* or downregulation of *grnd* rescue eye size, while downregulation of *veli* results in a small eye. Crosses at 25ºC; n=number of eyes (females); Kruskal-Wallis test (multiple comparisons).

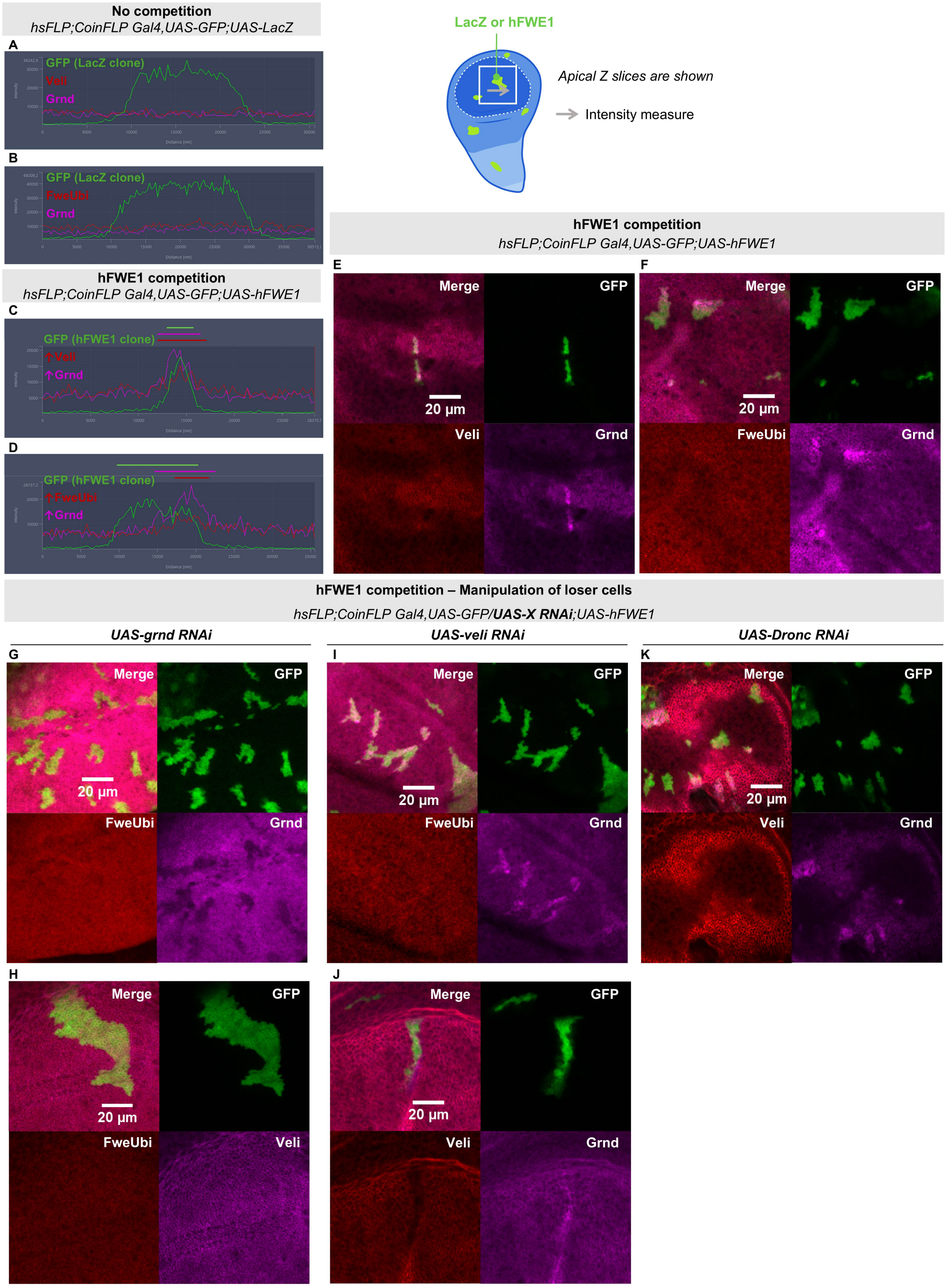


**Supplemental Fig.5 – Fwe-Grnd-Veli distribution in hFWE1-induced cell competition. A-D. Intensity profiles.** Intensity profiles of a clone in an apical slice are shown – A and B correspond to Fig.6 B and C, respectively; C and D correspond to Fig.6 D and E, respectively. **A-B.** In the absence of competition, Veli, Grnd and FweUbi are evenly distributed across cells. **C-D.** When hFWE1 cell competition is induced, Grnd can be detected in loser and winner cells at the interface. Veli and FweUbi accumulate mainly at the interface. **E-K.** Wing imaginal discs with GFP-labeled clones 48h and 72h after clone induction (*hsFLP;CoinFLP Gal4,UAS-GFP;UAS-X*) were imaged. A Z-plane slice of the apical membrane is shown at 20x magnification. **E-F.** hFWE1 loser cells accumulate Grnd apically and at borders. Veli and FweUbi do not show strong apical enrichment. **G-H.** Downregulation of *grnd* in loser clones leads to reduced Grnd signal, as expected, and FweUbi and Veli do not show strong alterations. **I-J.** Downregulation of *veli* in loser clones leads to reduced Veli signal, as expected. Strong Grnd enrichment in loser clones and in borders is detected, along with slight FweUbi accumulation apically in clones. **K.** Downregulation of *Dronc* still enables strong Grnd enrichment in loser clones and in borders, with some Veli enrichment also detected. The represented wing discs from E-K in this image correspond to the same discs shown in Figure 6 D-J.
